# Dynamic modelling of signalling pathways when ODEs are not feasible

**DOI:** 10.1101/2024.04.18.590024

**Authors:** Timo Rachel, Eva Brombacher, Svenja Wöhrle, Olaf Groß, Clemens Kreutz

**Affiliations:** Institute of Medical Biometry and Statistics, Medical Center, Faculty of Medicine, University of Freiburg, Stefan-Meier Str. 26, 79104, Freiburg, Germany; Institute of Physics, University of Freiburg, Hermann-Herder-Str. 3, 79104, Freiburg, Germany; CIBSS—Centre for Integrative Biological Signalling Studies, University of Freiburg, Schänzlestr. 18, 79104, Freiburg, Germany; Institute of Neuropathology, Faculty of Medicine, Medical Center, University of Freiburg, Breisacher Str. 64, 79106, Freiburg, Germany; Faculty of Biology, University of Freiburg, Schänzlestr. 1, 79104, Freiburg, Germany; Spemann Graduate School of Biology and Medicine (SGBM), University of Freiburg, Albertstr. 19A, 79104, Freiburg, Germany

**Keywords:** Mathematical modelling, Signalling pathways, Retarded transient function, Dose-dependency, Time-dependency, Predictions, Inflammasome activation, Tyrosine kinase inhibitors

## Abstract

**Motivation:** Mathematical modelling plays a crucial role in understanding inter- and intracellular signalling processes. Currently, ordinary differential equations (ODEs) are the predominant approach in systems biology for modelling such pathways. While ODE models offer mechanistic interpretability, they also suffer from limitations, including the need to consider all relevant compounds, resulting in large models difficult to handle numerically and requiring extensive data.

**Results:** In previous work, we introduced the *retarded transient function (RTF)* as an alternative method for modelling temporal responses of signalling pathways. Here, we extend the RTF approach to integrate concentration or dose-dependencies into the modelling of dynamics. With this advancement, RTF modelling now fully encompasses the application range of ordinary differential equation (ODE) models, which comprises predictions in both time and concentration domains. Moreover, characterizing dose-dependencies provides an intuitive way to investigate and characterize signalling differences between biological conditions or cell-types based on their response to stimulating inputs. To demonstrate the applicability of our extended approach, we employ data from time- and dose-dependent inflammasome activation in bone-marrow derived macrophages (BMDMs) treated with nigericin sodium salt. Our results show the effectiveness of the extended RTF approach as a generic framework for modelling dose-dependent kinetics in cellular signalling. The approach results in intuitively interpretable parameters that describe signal dynamics and enables predictive modelling of time- and dose-dependencies even if only individual cellular components are quantified.

**Availability:** The presented approach is available within the MATLAB-based *Data2Dynamics* modelling toolbox at https://github.com/Data2Dynamics and https://zenodo.org/records/14008247 and as R code at https://github.com/kreutz-lab/RTF.

## Introduction

Signalling networks play a crucial role in controlling cellular behavior, and deciphering these networks under different biological conditions is a fundamental goal of systems biology. One of the key approaches in this field involves quantifying the activation of cellular compounds and employing mathematical modelling to gain insights into signalling mechanisms and understand disease-causing malfunctions.

Traditionally, the prevailing modelling approach has been to translate known biochemical interactions into systems of ordinary differential equations (ODEs) using rate laws such as the law of mass action or enzyme kinetics. These mechanistic models, often referred to as rate-equation models, provide a mathematical representation of interactions of signalling compounds. In these models, each variable and parameter in the equations corresponds to a real biochemical entities and interaction strength within the pathway.

This approach offers mechanistic interpretability since each parameter corresponds to real biochemical entities, allowing for the precise assessment of compound contributions, integration of prior knowledge (e.g., typical concentrations or time scales), and predictions under conditions such as knockout or overexpression.

However, ODE-based mechanistic modelling also has certain limitations. The resulting models are nonlinear with respect to parameters, posing computational challenges such as numerical integration of high-dimensional stiff ODEs and dealing with local optima during parameter estimation. Additionally, statistical methods, which can account for the nonlinearity of the models need to be applied for model inference and uncertainty analysis. Another drawback is the necessity to include all relevant compounds, leading to large models that require substantial amounts of data to estimate the unknown model parameters.

To address these challenges, model reduction techniques have been proposed to decrease the model size by eliminating unnecessary components. However, model reduction limits mechanistic interpretability and can lead to biased reduction in model uncertainties.

As described in detail in [1], there are several alternative approaches for function estimation in the scientific literature. They fall into two main categories: nonparametric and parametric regression. Nonparametric methods like smoothing splines [2, 3], Gaussian processes [4], and kernel regression [5] do not require a predefined model structure, while parametric approaches, such as polynomial regression [6], non-linear regression techniques, and regression based on fractional polynomials [7, 8], rely on specified models. In systems biology, alternatives to ODE modelling have been developed to describe and investigate the dynamics of biochemical interactions. Gaussian processes are used for learning unknown differential functions [9] or inferring latent biochemical species [10]. Boolean models, which represent cellular signalling networks, can be transformed into continuous ODEs to serve as approximations. Additionally, dynamic Bayesian networks (DBNs) [11] offer ODE approximations with parallelized simulations on GPUs [12]. Logic-based ODEs have also been used to model cell signalling, combining logical rules with traditional ODE frameworks [13]. However, these alternative modelling approaches often result in uncommon dynamics and are not specifically tailored to cellular signalling processes.

The dynamics of cellular signalling processes show typically curve shapes [14]. We have previously proposed the retarded transient function (RTF) approach as an alternative modelling method specifically designed for dynamics commonly observed in ODE models derived by the rate-equation approach [1]. The non-mechanistic RTF approach incorporates characteristic features of signalling pathway responses, such as new steady states after stimulation, exhibiting monotone or at most one peak dynamics, delayed responses, e.g., for downstream compounds, and dynamics that are not orders of magnitude faster or slower than the measurement time scale. As the approach is describing transient dynamics, it is not suitable for several peaks or oscillations and therefore should not applied if such features play a prominent role in the dynamics.

In contrasts to mechanistic interpretability in ODE-based models, where parameters correspond to molecular interactions and rates, the RTF parameters capture the dynamics and can be interpreted intuitively in terms of amplitudes and time scales of the sustained and transient response components. Thereby they provide valuable biological insights in the systems dynamics.

We have shown that the RTF closely approximates the exact solutions of mechanistic ODEs in published pathway models and exhibits comparable predictive performance when fitted to simulated data and, thus, constitutes a valuable alternative modelling approach. The RTF is widely applicable, as shown recently by its use in the BayModTS workflow for analyzing diverse biological and clinical time course data [15].

However, so far, the RTF approach has been limited to describing and predicting time-dependencies while a major aim of systems modelling are predictions for varying stimulations or treatments which have not been feasible yet. In this work, we extend the RTF approach by incorporating dose-dependencies to model the effect of varying stimulation concentrations or doses with the help of Hill equations.

We demonstrate the ability of the novel dose-dependent RTF approach to effectively explain datasets, by applying it to measurements of inflammasome activation in bone-marrow derived macrophages (BMDMs) treated with nigericin. Additionally, we illustrate its utility in analyzing and interpreting the impact of genetic perturbations, such as the NEK7 knockout, on pathway responses. The presented modelling method is a valuable complementary tool for pathway modelling, particularly if only a few pathway components are measured or if the primary focus is on understanding the input-output behavior.

## Methodology

The RTF is a curve fitting approach which is based on three exponential functions and a non-linear transformation of the time axis. In the first section we recapitulate and illustrate the RTF as published previously [1] using a slightly different mathematical notation. Then, the RTF is generalized to multiple doses. finally, we illustrate the modelling and statistical testing of differences between different cell types or biological conditions.

### Single-dose RTF

The RTF defined as

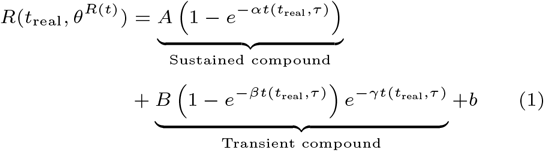

describes the time-dependency of the non-oscillating response of a cellular signalling compound by a sustained and a transient component. Both components have individual amplitudes *A, B* and time constants *α, β, γ*. In addition, an offset *b* and the magnitude *σ* of the measurement error of the data

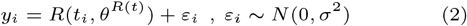

are estimated based on the measurements *y*_*i*_, *i* = 1, …, *N* at time points *t*_*i*_.

As the response of a pathway compound can be induced with some delay, a non-linear time transformation

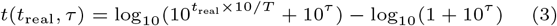

of the real experimental measurement times *t*_real_ is has been suggested [1]. The division by the range of the measurement times *T* makes the approach independent of time units. Time shift *τ* corresponds to the delay time and, using (3) as the argument of the exponential in (1) introduces a retarded response.

Overall, the time-dependent RTF comprises eight parameters *θ*^R(t)^ = {*A, B, α, β, γ, τ, b, σ*}, of which the first six parameters determine the dynamics. In this form, the time-dependent RTF only describes the temporal change of one signalling compound for one experimental condition, e.g., for one biological condition or for a single treatment dose. In the following, the approach will be generalized to multiple treatment doses. For the sake of simplicity, we will only use the term ‘dose’ in the following to describe the amount or concentration of an activating or suppressing signalling compound. To achieve a positive monotonic relationship between the dynamic parameters and multiple doses as introduced in the following section, we have parameterized the single-dose RTF using rate constants *α, β, γ*. In an application setting, the reciprocals of *α, β, γ* can be interpreted as time scales.

### Dose-dependent RTF

In biochemistry and pharmacology, Hill equations are often used to describe how a response depends on the dose of an activator, drug, or binding partner [16]. The Hill equation

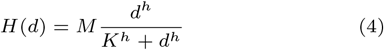

has three parameters: the maximum value *M*, the half-maximal quantity *K*, and the Hill coefficient *h*, which controls the sigmoidality. The half-maximal quantity *K* corresponds to the dose where half of the maximum activation is reached, and, in the context of pharamcokinetics, is often referred to as EC50. A visualisation of how these parameters effect the Hill equation are displayed in Tab. **??** in the supplements.

In the chosen parametrization, the response delay parameter *τ* is the only parameter with a negative monotonic relationship with increasing dose. Therefore, the corresponding Hill equation is modified to show a negative monotonic relationship with increasing doses:

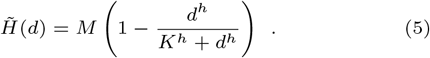

In order to obtain a dose-dependent formulation of the RTF, we use Hill equations to replace the six dynamic parameters of the time-dependent RTF *R*(*t, θ*^*R*(*t*)^) to describe how the kinetics depend on dose *d*:

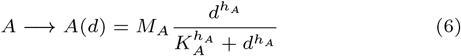

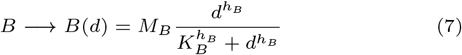

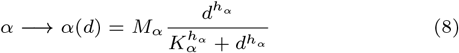

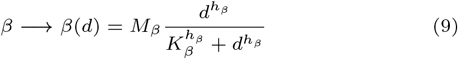

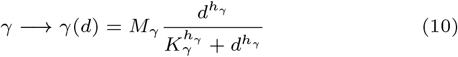

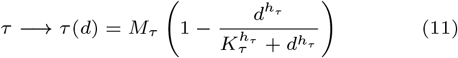

This is phenomenological description of the dose-dependency. For a motivation see section **??** in the supplements. Plugging these replacements into (1) results in the dose-dependent RTF *R*(*d, t, θ*^*R*(*t,d*)^) which has in its most general form 20 parameters

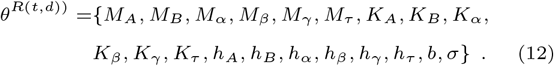

In case there is no dose-dependency, all half-maximal doses *K* are zero, resulting in all Hill equations becoming constant. In this case, the Hill coefficients *h* no longer have any influence and the dose-dependent RTF simplifies to the time-dependent RTF where the parameter values are given by the amplitudes *M*. In other words, the single-dose RTF represents the special case where *K*_*A*_ = *K*_*B*_ = *K*_*α*_ = *K*_*β*_ = *K*_*γ*_ = *K*_*τ*_ = 0 and *M*_*A*_ = *A, M*_*B*_ = *B, M*_*α*_ = *α, M*_*β*_ = *β, M*_*γ*_ = *γ, M*_*τ*_ = *τ* are constants.

Fig. 1 illustrates how the dose-dependencies of the parameters *A* and *τ* shown in (a) and (c) impact the corresponding dose-dependent RTFs. For most parameters of the RTF an increasing dose results in an stronger effect as exemplified for amplitude *A* of the sustained RTF part in panel (b). The time shift *τ* decreases with higher doses, prompting a faster response of the RTF Fig. 1 as shown in (d). Tab. **??** in the supplements contains similar figures for all parameters of the dose dependent RTF.

**Fig. 1.**
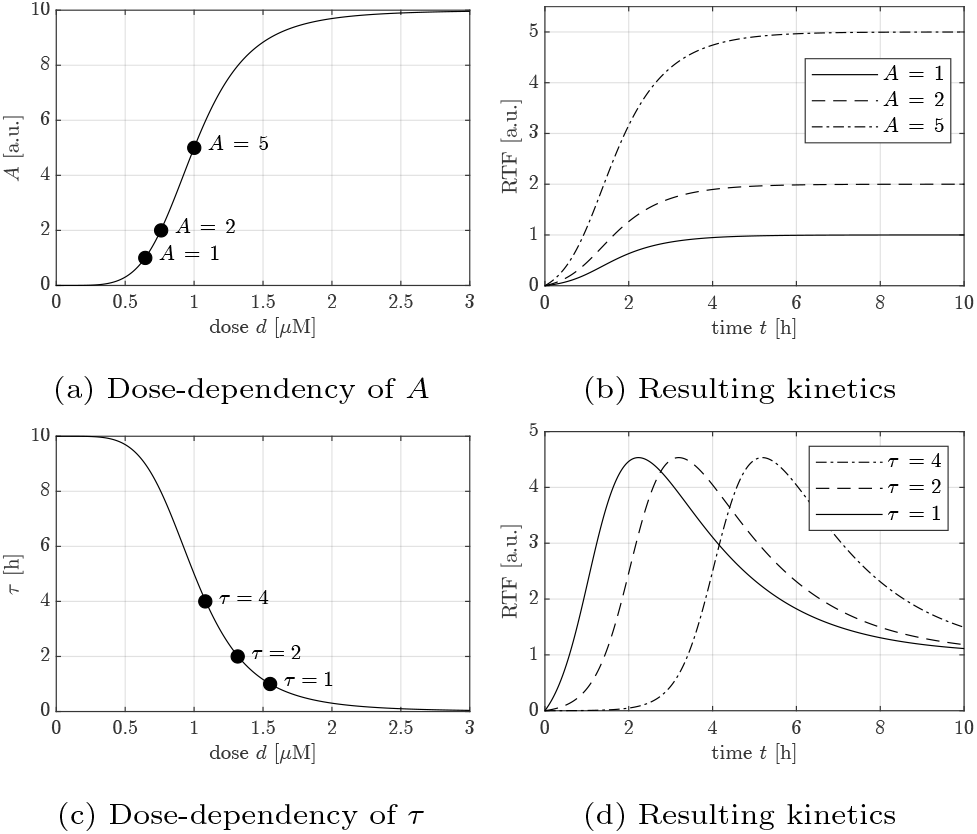
Illustrations of how dose-dependence of dynamic parameters translates into kinetics. See Tab. **??** in the supplements for a complete overview. **(a)** Hill curve showing the dose-dependency of amplitude *A*. **(b)** Resulting kinetics for the three values of *A* highlighted in (a). For better clarity, only the sustained part is displayed and the response saturates to the value of *A*. **(c)** The dose-dependency of the response time *τ* follows an decreasing Hill curve. **(d)** The three highlighted values of *τ* in (c) impact the time shift *τ* of the RTF response curves.

### Dose-dependent RTF for multiple conditions

Since in field of molecular biology typically several biological conditions are compared, e.g., different tissues, cell types, treatments, or genetically modified cells, we further generalize the dose-dependent RTF by introducing condition-dependent parameters

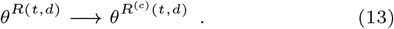

Here, the superscript index in parentheses, i.e., ^(*c*)^, refers to the considered biological conditions.

If dose-dependent time course datasets are measured for multiple biological conditions, interpreting differences from a reference condition is often of primary interest. To characterize differences in the RTF parameters over multiple conditions, we define a reference condition and introduce fold-changes, i.e., two conditions are parameterized with multiple fold-change parameters Δ by

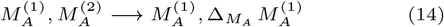

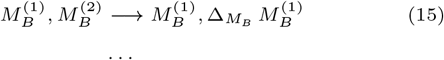

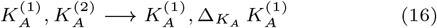

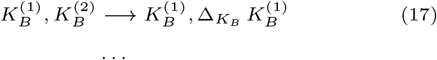

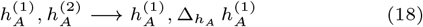

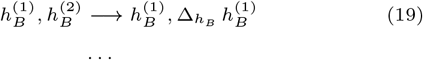

if condition (1) is considered as reference.

### Parameter estimation

The dose-dependent RTF can describe experimental data *y*(*t*) from time course data measurements across various doses and biological conditions, including those with replicates. For the jointly estimation of all of the model parameters in comparison to the data, the maximum likelihood approach is used, i.e., the estimated parameters

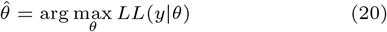

maximize the log-likelihood *LL*(*y*|*θ*).

To account for non-linearity during optimization, and especially to ensure the identification of the global minimum, we employ multi-start optimization [17]. The so-called *waterfall plot* visualization (see Fig. **??** in supplements) is used to check the reliability of the numerical optimization and to guarantee that the best optimum has been found repeatedly.

In the implementation chosen here, a single parameter *σ*, which quantifies the magnitude of the measurement errors, is jointly fitted with all other parameters. However, other error models, such as absolute and relative error models, or multiple *σ* parameters for groups of data points with different measurement errors, could be implemented if needed based on the data.

### Confidence intervals and significant condition-dependencies

Given that the retarded transient functions (RTFs) depend non-linearly on the parameters, and considering the usual limitations in data availability within the systems biology domain, it is crucial to avoid using local linear approximations for uncertainty quantification. Instead, uncertainty quantification methods that are specifically designed to handle non-linear systems should be used. This ensures the accurate and reliable estimation of uncertainties associated with RTF parameters and predictions. Thus, for the estimation of confidence intervals, we apply the profile likelihood approach [18], which has been proven to account for non-linearity and is one of the most frequently applied approaches for pathway models [19].

The profile likelihood provides a continuous evaluation of the likelihood ratio test (LRT) statistic for individual parameters [20]. In the context of systems biology, the profile likelihood is typically calculated on the log-scale because parameter are strictly positive and may vary over orders of magnitudes. By calculating the profile likelihood and setting a threshold which corresponds to a predetermined confidence level, the results can be interpreted in relation to the LRT. For instance, if the value Δ = 0 falls outside the 95% confidence interval, it implies a significant likelihood ratio test with a significance level of 5%. To statistically assess whether a fold-change parameter significantly differs from 1 (i.e., log-fold-change log(Δ) 0 for a given experimental condition), one can evaluate the profile likelihood for the log-fold-change parameter. The null hypothesis log(Δ) = 0 is rejected, if this value lies outside the respective confidence interval, i.e., the profile likelihood for log(Δ) = 0 is above the significance threshold.

### Parameter constraints and model reduction

In case data availability is limited, mathematical models with an excessive number of parameters can give rise to overfitting issues. To mitigate this issue in the context of time-dependent RTF modelling, we previously suggested to specify parameter bounds and perform model reduction [1].

The default parameter bounds were adapted to the dose-dependent RTF as shown in Tab. 1. The bounds for the maximum values *M*_*θ*_ match the ones for the corresponding parameters *θ*^*R*(*t*)^ proposed in [1]. The bounds proposed for *M*_*A*_ and *M*_*B*_ assume a model of up-regulation of the transient and sustained parts. If one of the amplitudes *A* or *B* is expected to be a down-regulation the respective sign can be changed or the bounds can be set to [−ub, 0] = [−2(max(*y*) − min(*y*)), 0]. *y* refers to the experimental time course data of all measured doses. If multiple conditions are compared, only the data of the reference condition is used for the definition of the boundaries. As the sustained and the transient part of the response are usually triggered together, *α* ≡ *γ* is assumed by default. For the Hill coefficient *h* and the half maximum quantity of the Hill equation *K* new default bounds are proposed here. And for all dose-dependent parameters {*A, B, α, β, γ, τ* } the same bound for *h* and *K* is used.

For dose-dependent RTF modelling, being applied both to single and multiple conditions, we recommend applying model reduction by a backward elimination procedure. This iterative approach facilitates repeated removal of parameters that are deemed unnecessary for adequately capturing the data. Furthermore, depending on the specific biological context, it may be desired to predefine parameters as being not dose-dependent, e.g., by setting the respective *K* to zero, based on prior knowledge. Additionally, not all parameters need to be condition-specific due to the underlying biological setting and it could be plausible to assume that certain parameters are shared among conditions.

## Results

To demonstrate its applicability, the extended RTF approach is applied to data quantifying NLRP3 inflammasome activation in bone-marrow derived macrophages (BMDMs) triggered by the bacterial toxin nigericin. An inflammasome is a multi-protein complex which initiates inflammatory responses and can induce cell death. During inflammasome activation interleukin-1β (IL-1β) is released. An enzyme-linked immunosorbent assay (ELISA) was used for detecting and quantifying IL-1β. Measurements were taken for a fixed set of time points for different treatment doses and different biological conditions.

The experiment was conducted for two experimental conditions: with wild-type cells and with NEK7 knockout cells.

In a fist step the time-dependent RTF, introduced in [1], is fitted for each treatment dose separately to the dynamics of the data from wild-type cells. Then, the dose-dependent RTF introduced in this manuscript is fitted to the wild-type data for all treatment doses in a joint fit, and the results are compared with the individual fits. Finally, the dose-dependent RTF for multiple conditions is fitted to both wild-type and knockout data to identify significant differences of the dose-dependent kinetics in knockout cells. For the parameter estimation of all models the bounds proposed in Tab.1 were used. They are shown in Tab. **??** to **??** in the supplements.

**Table 1.** Default parameter bounds and suggested initial values for parameter optimization.

| Parameter | Description | Lower bound (lb) | Upper bound (ub) | Default initial guess |
| --- | --- | --- | --- | --- |
| $M_A^*, M_B^*$ | Maximum value for Amplitude $A, B$ | 0 | $2(\max(y) - \min(y))^{**}$ | $0.1 \text{ lb} + 0.9 \text{ ub}$ |
| $M_\alpha, M_\gamma$ | Maximum value for rate constants $\alpha, \gamma$ | $1/(2(\max(t') - \min(t')))$ | $2/\min_i(t'_{i+1} - t'_i)$ | $0.5 \text{ lb} + 0.5 \text{ ub}$ |
| $M_\beta^{**}$ | Maximum value for rate constant $\beta$ | $M_\alpha$ | $M_\alpha$ | $M_\alpha$ |
| $M_\tau$ | Maximum value for time constant $\tau$ | $-(\max(t') - \min(t'))/5$ | $(\max(t') - \min(t'))/2$ | $-(\max(t') - \min(t'))/10$ |
| $h^{****}$ | Hill coefficient | 1 | 10 | $0.5 \text{ lb} + 0.5 \text{ ub}$ |
| $K^{****}$ | Half maximum quantity | $\min(\text{dose} > 0)/10$ | $\max(\text{dose}) \cdot 10$ | $0.5 \text{ lb} + 0.5 \text{ ub}$ |
| $b$ | Data offset | $\min(y)^{**}$ | $\max(y)^{**}$ | $0.5 \text{ lb} + 0.5 \text{ ub}$ |
| $\sigma$ | Measurement error of data | $\min(\text{ub}, \max(10^{-10}, (\max(y) - \min(y))/10^4))^{**}$ | $\max(10^{-10}, \text{SD}(y))^{**}$ | $0.5 \text{ lb} + 0.5 \text{ ub}$ |

\*For down-regulation switch and change sign: $\text{lb} = -2(\max(y) - \min(y))$ , $\text{ub} = 0$ .
\*\* $y$ denotes the data that is used for the parameter estimation of the model. For the comparison of multiple conditions only the data from the reference condition is used.
\*\*\*Since the sustained and the transient part of the response are usually triggered together, $\alpha = \gamma$ is assumed by default.
\*\*\*\*The suggested boundaries for the Hill coefficient and the half maximum quantity are identical for all dose-dependent parameters of the RTF.

### Modelling dose-dependent dynamics

In the first step the traditional single-dose RTF (1) was fitted for each treatment dose separately to the data of the wild-type cells. Based on prior biological knowledge we set *B* = 0 resulting in the RTF having no transient part. The remaining parameters are *θ*^R(t)^ = {*A, α, τ, b, σ*}. For each of the 7 treatment doses we get individual estimates for the model parameters. In total, the single-dose RTF requires 35 parameters over all 7 doses. These individual fits are based on and do not allow for predictions for unobserved doses. The resulting fits are shown in Fig. 2 (dashed lines).

**Fig. 2.**
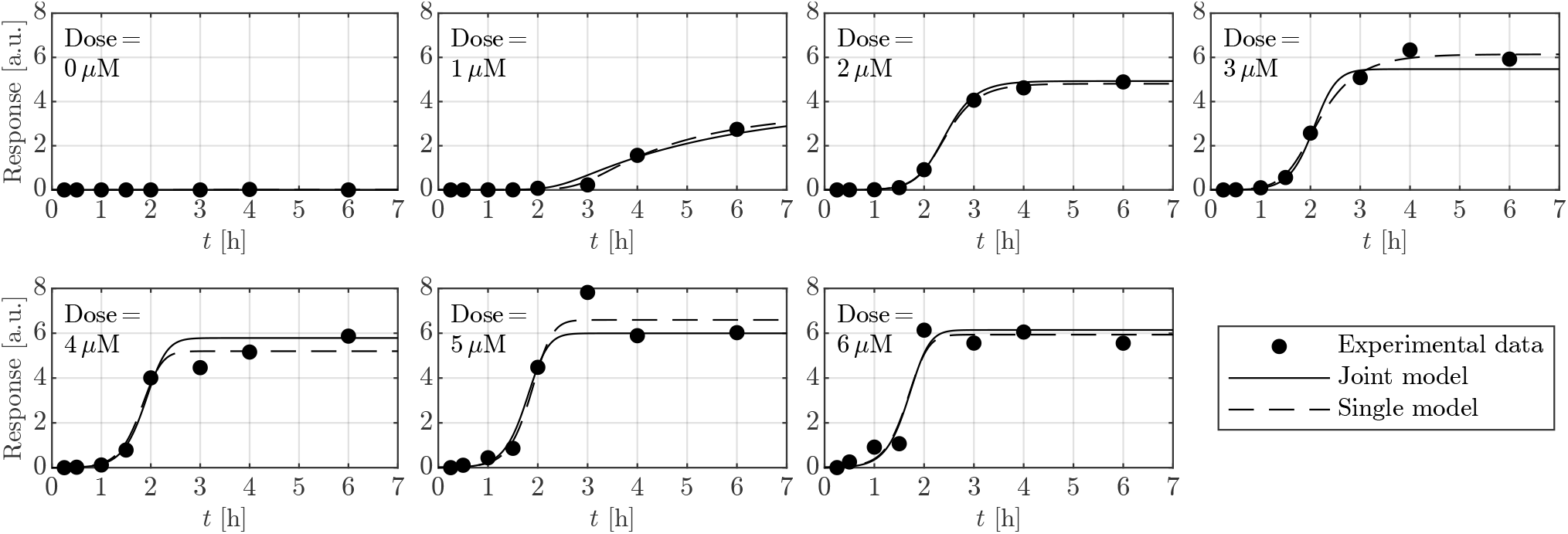
Comparison of fitting individual functions to each dose (single-dose RTF approach) with a joint fit of the dose-dependent joint model on the same experimental data. In contrast to the 35 parameters of the individual fits, the joint model requires only 11 parameters to describe the data and enables predictions for unobserved doses.

Next the dose-dependent RTF is used to model dynamics of the wild-type cells by jointly fitting all treatment doses simultaneously. For this purpose, the parameters *A, α, τ* are replaced with the Hill equations (6), (8) and (11) in (1) as described above. As before, we presumed that there is no transient part of the response and set the amplitude *B* to zero. In contrast to the in total 35 parameters that are needed to model the RTF for each dose individually, only 11 parameters are needed for the dose-dependent joint model independent of the number of doses: {*M*_*A*_, *M*_*α*_, *M*_*τ*_, *K*_*A*_, *K*_*α*_, *K*_*τ*_, *h*_*A*_, *h*_*α*_, *h*_*τ*_, *b, σ*}.

For each of the dose-dependent parameters *A, α*, and *τ* there are three Hill parameters: *M, K*, and *h*. Additionally, there the are three parameters *b* and *σ* that are independent of the dose. Although the dose-dependent RTF model requires fewer parameters and enforces that the dose-dependencies of the RTF parameters follow Hill functions, the fit to the data shows no visible loss in explaining the data (Fig. 2). To compare the dose-dependent RTF fit with the individually fitted RTFs, we used the fixed *σ* as fitted by the dose-dependent RTF for both models. This results in, *χ*^2^ = 56.0 for the dose-dependent RTF with 10 fitted parameters, and *χ*^2^ = 65.5 for the individually fitted RTFs to each of the 7 doses for 30 parameters in total. A likelihood-ratio test yields a p-value close to 1, indicating that there is no significant difference between the quality the dose-dependent RTF compared to the individual fits. As they have a larger amount of parameters the individual RTFs are over-parameterized compared to the dose-dependent RTF. Furthermore, the dose-dependent RTF ensures a monotonic increase of the response for all doses, which is not guaranteed in the individual fits (see Fig. 2 dashed lines for 4, 5, 6 µM). Despite this, the dose-dependent RTF describes dynamic responses to varying doses, enabling predictions of system dynamics for unmeasured treatment regimes.

### Modelling condition-dependencies

To demonstrate the applicability of the dose-dependent RTF across various conditions, it is applied to both wild-type cell data and the data obtained from the cells with a NEK7 knockout (Fig. 3). The corresponding joint fits are shown in Fig. 3. The wild-type serves as a reference condition and is augmented with fold-changes Δ for each parameter ((14) to (19)). The likelihood profiles of these fold-change parameters Δ are shown in Fig. 4, where the dashed line shows the 95% confidence level, with the *x*-axis displayed in logarithmic scale.

**Fig. 3.**
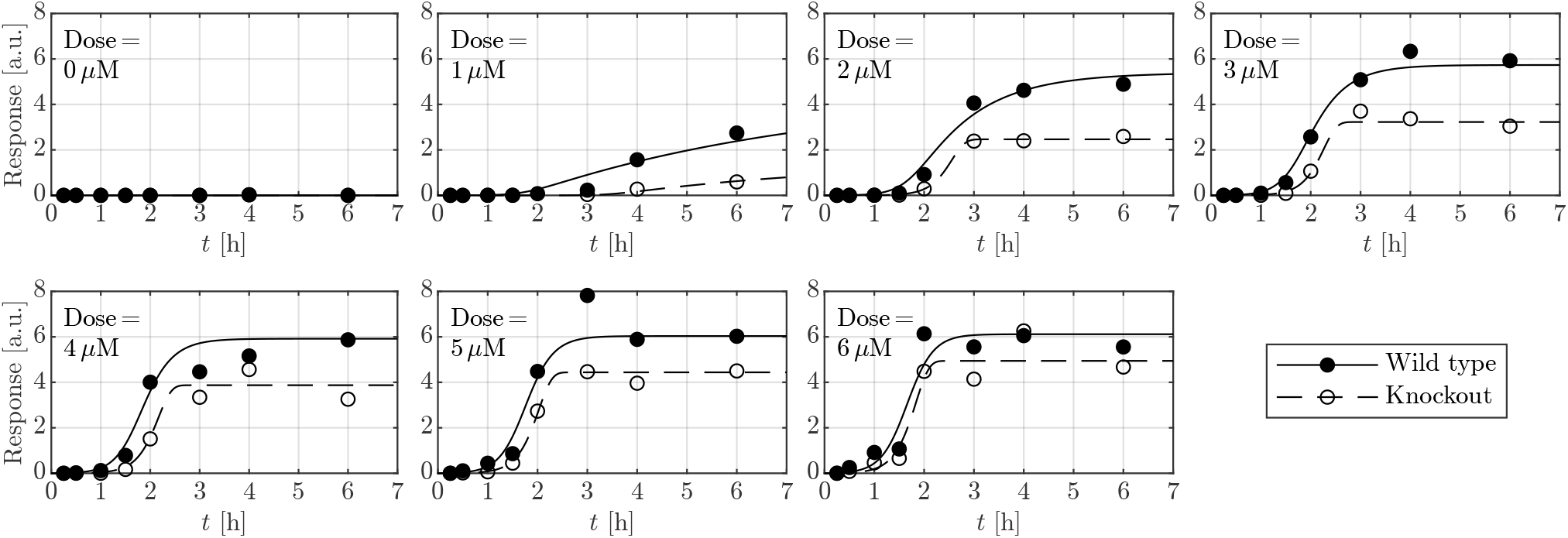
The condition-specific dose-dependent RTF can describe the data generated over two different conditions (wild-type cells and NEK7 knockout cells). Furthermore, fold-changes Δ can be estimated and tested for significance.

**Fig. 4.**
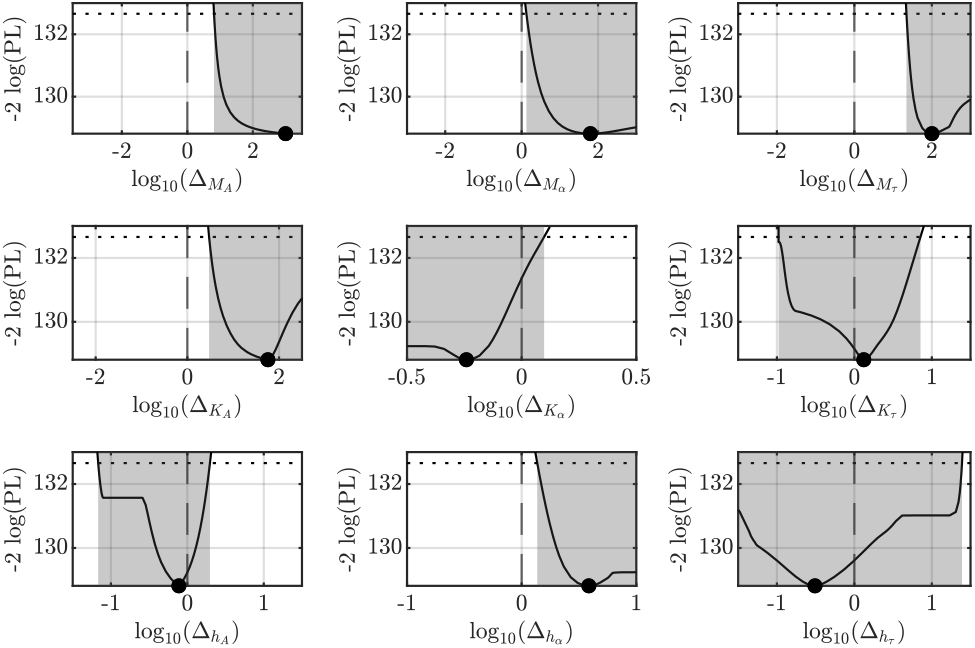
The profile likelihood of the fold-change parameters Δ can be used to test, which parameters are significantly different in the knockout condition. The intersection points between the dotted threshold and the profile likelihood indicate the margins of the 95% confidence interval (CI) highlighted by gray coloring. If the 0 on the logarithmic scale (corresponding to 1 on the linear scale) is outside the 95% CI the likelihood ratio test is significant to a 5% significance level, indicating significantly different characteristics of the dose-dependency between both conditions.

For fold-changes of *M*_*A*_, *M*_*α*_, *M*_*τ*_, *K*_*A*_, and *h*_*α*_ there is a significant change in NEK7 knockout cells indicated by fold-factors significantly different from one, i.e. the 95% confidence interval (CI) for the log-fold-changes do not cover zero. Conversely, for the parameters *K*_*α*_, *K*_*τ*_, *h*_*A*_, and *h*_*τ*_ the log-fold-change Δ = 0 lies within the 95% CI. As these factors do not significantly differ from 1 the corresponding parameters exhibit no condition-dependency. Therefore, a model reduction procedure could be applied by iteratively eliminating the condition-dependencies of these parameters. However, as with any model reduction procedures, such backward elimination might result in different solutions depending on the specific order in which parameters are removed.

## Conclusion

The development of a comprehensive toolbox of methods applicable and tailored to a broad range of applications and research questions is crucial for performing effective modelling in systems biology. This manuscript proposes a novel approach for dynamic modelling of cellular signalling processes, especially if ODEs are not feasible, e.g., due to computational problems arising from large model sizes or a lack of sufficient experimental data for all relevant compounds in a mechanistic model. Here, we extended the retarded transient function (RTF) to describe dose-dependencies typically observed in the context of cellular signalling. The approach’s applicability and efficacy is demonstrated at the example of the analysis of time- and dose-dependent measurements of inflammasome activation.

The presented dose-dependent modelling can be applied to individual signalling compounds to capture and explain experimental observations. Notably, the dose-dependent RTF requires fewer parameters than the single-dose RTF approach if applied to multi-dose experiments and substantially fewer then those needed for ODE modelling.

In contrast to mechanistic ODE models, individual observables can be modelled without the need for integrating all relevant compounds of a signalling process as required for mechanistic models.

Since the dose-dependent RTF model describes dynamic responses as a function of varying doses, it also enables predictions of system dynamics for unmeasured treatment doses. It also facilitates fitting of input output behavior of a system of interest in a less elaborate manner by a phenomenological, non-mechanistic mathematical description. Moreover, it offers novel opportunities to link mathematical models of related signalling processes, e.g. for multi-scale models.

Additionally, the intuitive interpretation of its parameters, e.g. as amplitudes, delays, and time rates, supports the characterization and understanding of signalling dynamics.

Furthermore, this approach enables statistically rigorous analysis of differences in the responses of biological pathways. This not only deepens our insight into cell signalling but also provides a structured framework for testing hypotheses and comparing various biological conditions such as wild-type and knockout.

In particular, when the amount of available data is limited, assumptions must be made for mathematical modelling in order to balance the trade-off between flexibility and overfitting. Therefore, the dose-dependent RTF approach has certain application limits, like any other modelling approach. As also discussed in [1], the single-dose RTF is tailored to non-oscillatory response kinetics, as typically observed as the initial response in ODE systems resulting from the rate equation approach. Therefore, responses with more than one peak or oscillations are not accurately described by the RTF but only effectively as an average over any oscillations that may occur. As shown in [1], the RTF is, nevertheless, very well suited for the vast majority of pathway models, so that the RTF can be applied for a broad range of applications.

In summary, the introduced dose-dependent RTF serves as a promising and interpretable complementary modelling approach for the investigation of cellular signalling processes. Its ability of accurately explaining data, providing intuitive insights, supporting predictions of time- and dose-dependencies, and facilitating statistical testing renders it a valuable addition to the repertoire of systems biology modelling techniques benefiting the understanding of dynamic signalling processes when ODEs are hardly feasible.

## Author contributions statement

O.G. and S.W. conceived and conducted the experiments, C.K. developed the theoretical formalism, T.R. and E.B. implemented the approach, T.R. performed the simulations and analysed the results supervised by C.K., C.K., T.R. and E.B. wrote and reviewed the manuscript.

## Acknowledgment

This study was funded by the Deutsche Forschungsgemeinschaft (DFG, German Research Foundation) under Germany’s Excellence Strategy (CIBSS—EXC-2189—Project ID 390939984) and the University of Freiburg in the funding program Open Access Publishing.

