## supplementary material for "Dynamic modelling of signalling pathways when ODEs are not feasible"

### Contents

**S1 Waterfall Plot** **II**

**S2 Motivation of Hill functions for describing dose dependencies** **III**

**S3 Illustration of Parameters** **IV**
S3.1 Single-Dose RTF . . . . . IV
S3.2 Hill Function . . . . . VI
S3.3 Dose-Dependent RTF . . . . . VII

**S4 Boundaries and Confidence Intervals for all Scenarios** **XII**
S4.1 Modelling doses individually . . . . . XII
S4.1.1 Dose 1 . . . . . XII
S4.1.2 Dose 2 . . . . . XII
S4.1.3 Dose 3 . . . . . XII
S4.1.4 Dose 4 . . . . . XII
S4.1.5 Dose 5 . . . . . XIII
S4.1.6 Dose 6 . . . . . XIII
S4.1.7 Dose 7 . . . . . XIII
S4.2 Modelling dose-dependent dynamics . . . . . XIV
S4.3 Modelling condition-dependencies . . . . . XV

### S1 Waterfall Plot

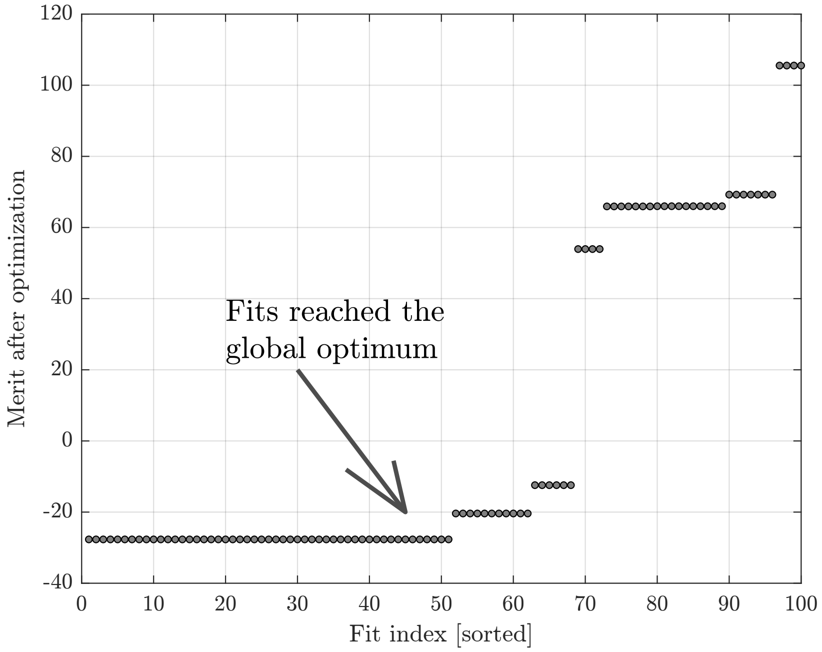


Figure S1: Example of a waterfall plot used to identify the global optimum of the parameters for the dose-dependent RTF of the fits shown in Fig. **??**. The parameter optimization initializes $n = 100$ times, each time starting with a random parameter vector within specified boundaries. In the waterfall plot, the objective function value of each of the $n$ optimization runs corresponds to the $y$-axis value, ordered from smallest to largest objective function value. This results in a visualization of different merit levels, each level corresponding to a local optimum. The lowest level is assumed to be the global optimum.

### S2 Motivation of Hill functions for describing dose dependencies

Hill functions provide a convenient and intuitive way to describe how the steady-state response of a biochemical system depends on the concentration of an input (like a drug or substrate). This relationship is rooted in the underlying ODEs that describe the system's dynamics as shown in the following. The formation of a complex $C$ by binding of an activator $A$ to a substrate $S$ translates to

$$[\dot{C}] = k_f[A][S] - k_b[C] \tag{1}$$

using the law of mass action for the forward and backward reaction with two rate constants $k_f$ and $k_b$. For the steady state $[\dot{C}] = 0$, the equation becomes

$$[C] = \frac{k_f}{k_b}[A][S] \ . \tag{2}$$

By substituting $A = A_{\text{total}} - C$ the equation yields

$$[C] = \frac{k_f}{k_b}(A_{\text{total}} - C)[S] \tag{3}$$

and after solving for $C$ we obtain

$$\begin{aligned}
[C] &= \frac{k_f}{1 + \frac{k_f}{k_b}}[S] && (4)\\
&= \frac{\frac{k_b}{k_f}k_f[S]}{\frac{k_b}{k_f} + \frac{k_b}{k_f}\frac{k_f}{k_b}[S]} && (5)\\
&= \frac{V_{\max}[S]}{K_D + [S]} && (6)
\end{aligned}$$

with $K_D := k_b/k_f$ and $V_{\max} = k_b$. This result corresponds to a Hill function

$$H([S]) = \frac{V_{\max}[S]^h}{K_D^h + [S]^h} \tag{7}$$

with Hill coefficient $h = 1$.

The steady state of the RTF is represented by the amplitude $A$ of the sustained component. In the dose-dependent formulation, the Hill function $A(d)$, which describes how the amplitude $A$ depends on the dose, follows the steady-state relationship typically derived from simple complex formation. More complex processes, as seen in biochemical systems, can also be described by Hill coefficients $h \neq 1$."

In more complex biochemical networks, also other parameters of the RTF can be described by Hill functions. Let's assume that the complex $C$ regulates another target $T$ via

$$[\dot{T}] = k_t[C][T] \ . \tag{8}$$

For the immediate effect on the target $T$, Taylor expansion of $[T](t)$ around time point $t = 0$ yields

$$\begin{aligned}
[\dot{T}](t) &\approx k_t[C](t)([T](t=0) + t[\dot{T}](t=0) + \dots) && (9)\\
&\approx k_t[C](t)[T](t=0) \ \text{for} \ t \approx 0 \ . && (10)
\end{aligned}$$

Thus, when $[C]$ has a dose-dependency that is described by a Hill function, the immediate effect on targets of $C$, described by rate $\alpha$ for the sustained component and $\beta$ for the transient component also have a Hill dose-dependency described by a Hill function $\alpha(d)$ and $\beta(d)$.

In more complex regulation networks and for downstream targets, this effect might be delayed and is accordingly described by the retarded RTF and the time shift parameter $\tau$. The dose-dependent RTF also has the flexibility to describe the dose-dependency of such a delayed response by the sigmoidal Hill function $\tau(d)$.

In strict mathematical terms, it cannot be shown that in complex signalling processes, all compounds can be described by Hill function of the RTF parameters. After all, the dose-dependent RTF approach is a phenomenological model.

### S3 Illustration of Parameters

#### S3.1 Single-Dose RTF

As specified in Equations (**??**) and (**??**), the single-dose RTF is given by

$$R(t_{\text{real}}, \theta^{R(t)}) = A\left(1 - e^{-\alpha t(t_{\text{real}},\tau)}\right) + B\left(1 - e^{-\beta t(t_{\text{real}},\tau)}\right) e^{-\gamma t(t_{\text{real}},\tau)} + b \ ,$$

where

$$t(t_{\text{real}}, \tau) = \log_{10}(10^{t_{\text{real}} \times 10/T} + 10^{\tau}) - \log_{10}(1 + 10^{\tau}) \ .$$

Table S1 shows how the single-dose RTF is affected by its parameters.

Table S1: Parameters of the single-dose Retarded Transient Function (RTF) and their effects. For each described parameter, an example plot is provided, where the respective parameter is varied from low (blue) to high (red). Next to the example plots the used parameter values are listed.

| Parameter | Explanation | Illustration of the Effect on the RTF |
|---|---|---|
| $A$ | Amplitude of the sustained response. For clarity, in this example, $B = 0$ i.e., only the sustained part remains. | 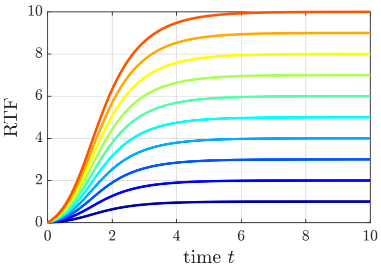 $A = 1$ to $10$<br>$B = 0$<br>$\alpha = 1$<br>$\tau = 1$<br>$b = 0$ |
| $B$ | Amplitude of the transient response. For illustration purposes, in this example is $A = 0$, i.e., only the transient part remains. Note that $B$ is reached only if $\beta \gg \gamma$ since the transient response approaches amplitude $B$ with rate $\beta$ and, at the same time, decays with rate $\gamma$. | 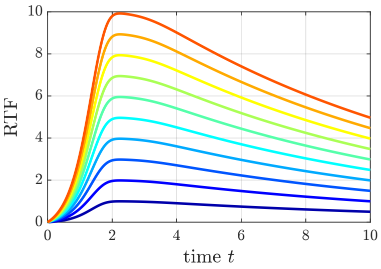 $A = 0$<br>$B = 1$ to $10$<br>$\beta = 100$<br>$\gamma = 0.1$<br>$\tau = 3$<br>$b = 0$ |
| $\alpha$ | The rate constant of the sustained response $\alpha$ controls how fast amplitude $A$ of the sustained part is approached and, thus, corresponds to the steepness of slope of the sustained part. In this example $B = 0$, i.e., only the sustained part remains. | 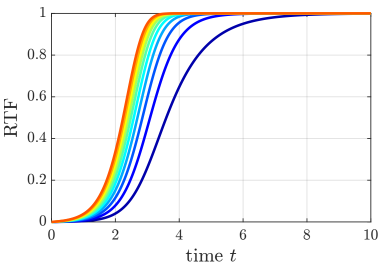 $A = 1$<br>$B = 0$<br>$\alpha = 1$ to $10$<br>$\tau = 3$<br>$b = 0$ |

| | | |
|---|---|---|
| $\beta$ | The rate constant of the transient response $\beta$ describes how fast the transient part approaches amplitude $B$. As $B$ is only reached if $\beta \gg \gamma$, $\beta$ and $\gamma$ indirectly impact the strength of the transient part. | 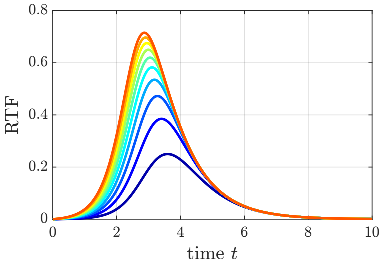<br>$A = 0$<br>$B = 1$<br>$\beta = 1$ to $10$<br>$\gamma = 1$<br>$\tau = 3$<br>$b = 0$ |
| $\gamma$ | The rate constant of the transient decay $\gamma$ escribes how fast the transient part decays. Increasing $\gamma$ also reduced the maximum value of the RTF. | 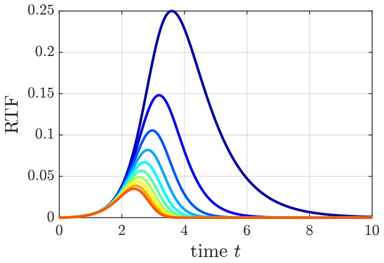<br>$A = 0$<br>$B = 1$<br>$\beta = 1$<br>$\gamma = 1$ to $10$<br>$\tau = 3$<br>$b = 0$ |
| $\tau$ | The response time $\tau$ describes a shift along the $x$-axis, i.e., can account for delayed responses. | 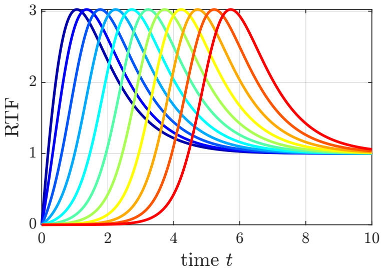<br>$A = 1$<br>$B = 10$<br>$\alpha = 1$<br>$\beta = 1$<br>$\gamma = 1$<br>$\tau = 0$ to $5$<br>$b = 0$ |
| $b$ | The data offset $b$ can introduce a constant vertical shift. | 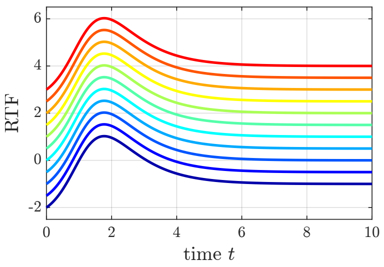<br>$A = 1$<br>$B = 10$<br>$\alpha = 1$<br>$\beta = 1$<br>$\gamma = 1$<br>$\tau = 1$<br>$b = -2$ to $3$ |

### S3.2 Hill Function

As specified in Equation (**??**), the Hill equation is given by

$$H(d) = M \frac{d^h}{K^h + d^h} \; .$$

Table S2 shows how the Hill equation is affected by its parameters.

Table S2: Parameters of the Hill equation and their effect. For each described parameter, an example plot is provided, where the respective parameter is varied from low (blue) to high (red). Next to the example plots the used parameter values are listed.

| Parameter | Explanation | Illustration of the Effect on the Hill Function |
|---|---|---|
| $M$ | The maximum value $M$ of the Hill function is asymptotically approached for large doses $d$. | 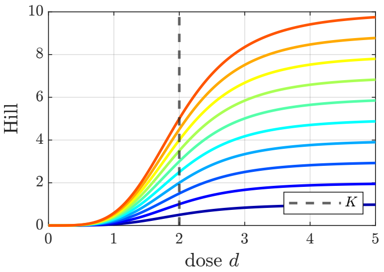  $M = 1$ to $10$, $K = 2$, $h = 4$ |
| $K$ | The half maximum quantity is the dose that produces 50% of the maximal value ($\frac{M}{2}$), which relates to the steepness of the curve. | 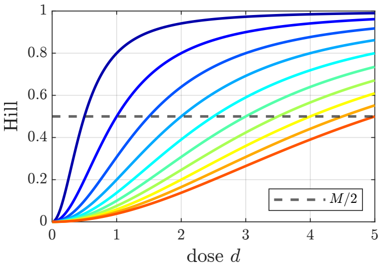  $M = 1$, $K = 0.5$ to $5$, $h = 2$ |
| $h$ | The Hill coefficient $h$ defines the sigmoidality of the curve. When varying $h$, all resulting curves have a common intersection at the point $(K, M/2)$. For doses smaller than $K$ curves with smaller $h$ values increase faster, while for doses greater than $K$ curves with smaller $h$ values increase slower. | 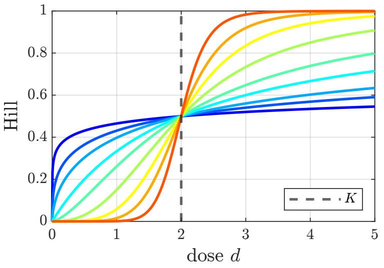  $M = 1$, $K = 2$, $h = 0.2$ to $10$ |

### S3.3 Dose-Dependent RTF

As specified in Equations (**??**), and (**??**) to (**??**) the dose-dependent RTF is given by

$$R(t_{\text{real}}, \theta^{R(t)}) = M_A \frac{d^{h_A}}{K_A^{h_A} + d^{h_A}} \left(1 - e^{-M_\alpha \frac{d^{h_\alpha}}{K_\alpha^{h_\alpha} + d^{h_\alpha}} t(t_{\text{real}}, \tau)}\right)$$
$$+ M_B \frac{d^{h_B}}{K_B^{h_B} + d^{h_B}} \left(1 - e^{-M_\beta \frac{d^{h_\beta}}{K_\beta^{h_\beta} + d^{h_\beta}} t(t_{\text{real}}, \tau)}\right) e^{-M_\gamma \frac{d^{h_\gamma}}{K_\gamma^{h_\gamma} + d^{h_\gamma}} t(t_{\text{real}}, \tau)} + b \ ,$$

with

$$t(t_{\text{real}}, \tau) = \log_{10}\left(10^{t_{\text{real}} \times 10/T} + 10^{M_\tau \left(1 - \frac{d^{h_\tau}}{K_\tau^{h_\tau} + d^{h_\tau}}\right)}\right) - \log_{10}\left(1 + 10^{M_\tau \left(1 - \frac{d^{h_\tau}}{K_\tau^{h_\tau} + d^{h_\tau}}\right)}\right) \ .$$

Table S3 shows how the dose-dependent RTF is affected by its parameters.

Table S3: Parameters of the dose-dependent Retarded Transient Function (RTF) and their corresponding Hill functions. In each row, one parameter is varied. For selected doses, the corresponding RTFs are plotted, where the colors (blue, red, or yellow) of the RTF plots are equal to the corresponding dots in the Hill graphs.

| Parameter | Explanation | Effect on Hill function | Effect on the RTF |
|---|---|---|---|
| $M_A$ | Maximum value or saturation of the Hill function for amplitude $A$ of the sustained RTF part, reached for high doses. In this example $B = 0$, i.e., only the sustained part is set to 0. | 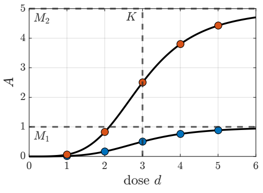 $M = 1, 5$<br>$K = 3$<br>$h = 4$ | 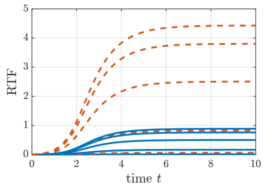 $B = 0$<br>$\alpha = 1$<br>$\tau = 2$<br>$b = 0$ |
| $K_A$ | The so-called half maximum quantity $K_A$ of the Hill function specifies the dose for which the amplitude $A$ of the sustained RTF part is 50% of the maximal amplitude $M_A$. | 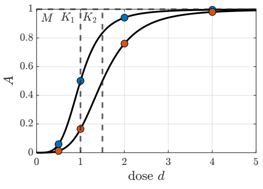 $M = 1$<br>$K = 1, 1.5$<br>$h = 4$ | 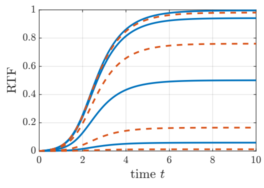 $B = 0$<br>$\alpha = 1$<br>$\tau = 2$<br>$b = 0$ |
| $h_A$ | Hill coefficient $h$ of the Hill function for amplitude $A$ of the transient RTF part. Increasing $h_A$ enhances the sigmoidality of the Hill curve, making it steeper. This steepening effect leads to a sudden increase of amplitude $A$ of the transient RTF part, particularly for doses near $K$. | 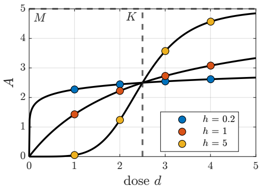 $M = 5$<br>$K = 2.5$<br>$h =$<br>$0.2, 1, 5$ | 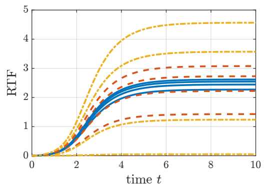 $A = 0.5$<br>$\alpha = 10$<br>$\beta = 10$<br>$\gamma = 1$<br>$\tau = 2$<br>$b = 0$ |

| | | | | | |
|---|---|---|---|---|---|
| $M_B$ | Maximum value of the amplitude $B$ of the transient RTF part, reached for high doses. Note that $B$ is typically not reached as the signal decays with rate $\gamma$. In the example plot the transient part has nearly vanished for $t = 6$ and the response is determined by the amplitude of the sustained part $A$. | 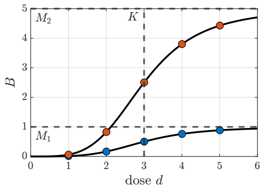 | $M = 1, 5$<br>$K = 3$<br>$h = 4$ | 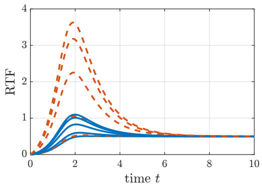 | $A = 0.5$<br>$\alpha = 10$<br>$\beta = 10$<br>$\gamma = 1$<br>$\tau = 2$<br>$b = 0$ |
| $K_B$ | The half maximum quantity $K_B$ of the Hill function specifies the dose for which the amplitude $B$ of the transient RTF part is 50% of the maximal amplitude $M_B$. | 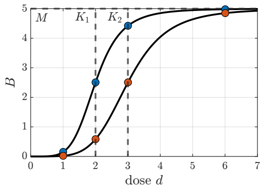 | $M = 5$<br>$K = 2, 3$<br>$h = 5$ | 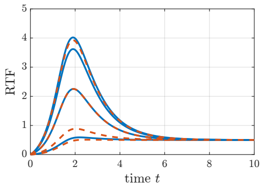 | $A = 0.5$<br>$\alpha = 10$<br>$\beta = 10$<br>$\gamma = 1$<br>$\tau = 2$<br>$b = 0$ |
| $h_B$ | Hill coefficient $h$ of the Hill function for amplitude $B$ of the transient RTF part. Increasing $h_B$ enhances the sigmoidality of the Hill curve, making it steeper. This steepening effect leads to a sudden increase of amplitude $B$ of the transient RTF part, particularly for doses near $K$. | 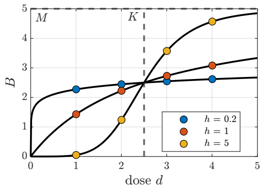 | $M = 5$<br>$K = 2.5$<br>$h =$<br>$0.2, 1, 5$ | 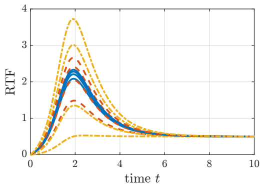 | $A = 0.5$<br>$\alpha = 10$<br>$\beta = 10$<br>$\gamma = 1$<br>$\tau = 2$<br>$b = 0$ |
| $M_\alpha$ | Maximum value for the rate constant $\alpha$ of the sustained RTF part, reached for high doses. For the example plot, the transient part is set to 0. | 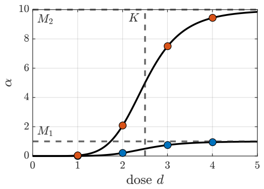 | $M = 1, 10$<br>$K = 2.5$<br>$h = 6$ | 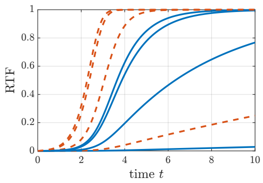 | $A = 1$<br>$B = 0$<br>$\tau = 3$<br>$b = 0$ |

| | | | |
|---|---|---|---|
| $K_\alpha$ | The half maximum quantity $K_\alpha$ of the Hill function specifies the dose for which the rate constant $\alpha$ of the sustained RTF part is 50% of the maximal rate $M_\alpha$. Roughly speaking, $K_\alpha$ indicates the dose at which the "velocity" is 50% of the maximum velocity of the sustained part. | 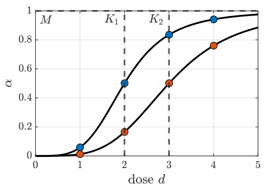 $M = 1$<br>$K = 2, 3$<br>$h = 4$ | 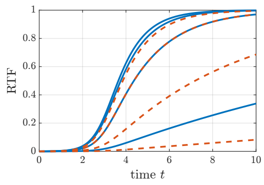 $A = 1$<br>$B = 0$<br>$\tau = 3$<br>$b = 0$ |
| $h_\alpha$ | The Hill coefficient $h_\alpha$ controls how non-linear the rate constant $\alpha$ depends on the dose. Increasing $h_\alpha$ enhances the sigmoidality of the Hill curve, making it steeper. Large values of $h$ leads to a sudden increase of the "velocity" $\alpha$ for doses close to $K_\alpha$. | 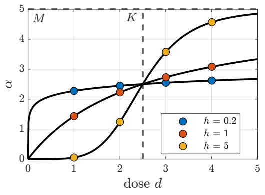 $M = 5$<br>$K = 2.5$<br>$h =$<br>$0.2, 1, 5$ | 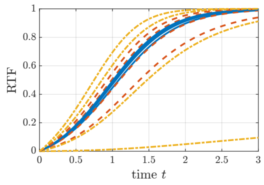 $A = 1$<br>$B = 0$<br>$\tau = 3$<br>$b = 0$ |
| $M_\beta$ | Maximum value for the rate constant $\beta$ of the sustained RTF part, reached for high doses. For the example plot, the sustained part is set to 0. | 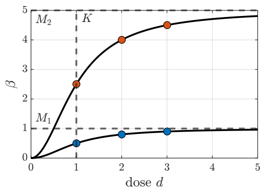 $M = 1, 5$<br>$K = 1$<br>$h = 2$ | 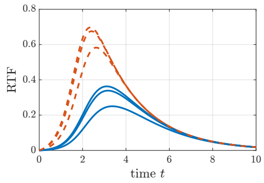 $A = 0$<br>$B = 1$<br>$\gamma = 0.5$<br>$\tau = 2$<br>$b = 0$ |
| $K_\beta$ | The half maximum quantity $K_\beta$ of the Hill function specifies the dose for which the rate constant $\beta$ of the transient RTF part is 50% of the maximal rate $M_\beta$. Roughly speaking, $K_\beta$ indicates the dose at which the "velocity" is 50% of the maximum velocity of the increase of the transient part. | 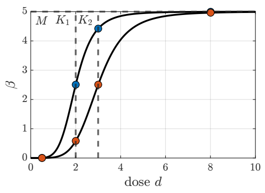 $M = 5$<br>$K = 2, 3$<br>$h = 5$ | 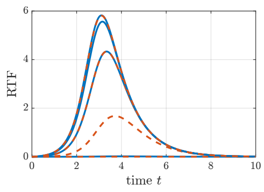 $A = 0$<br>$B = 10$<br>$\gamma = 1$<br>$\tau = 3$<br>$b = 0$ |

<table>
<tr>
<td>$h_\beta$</td>
<td>The Hill coefficient $h_\beta$ controls how non-linear the rate constant $\beta$ depends on the dose. Increasing $h_\beta$ enhances the sigmoidality of the Hill curve, making it steeper. Large values of $h$ leads to a sudden increase of the "velocity" $\beta$ for doses close to $K_\beta$.</td>
<td>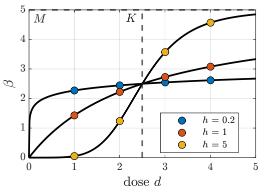<br>$M = 5$<br>$K = 2.5$<br>$h =$<br>$0.2, 1, 5$</td>
<td>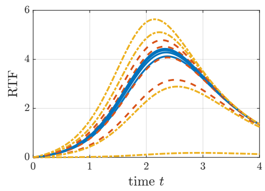<br>$A = 0$<br>$B = 10$<br>$\gamma = 1$<br>$\tau = 2$<br>$b = 0$</td>
</tr>
<tr>
<td>$M_\gamma$</td>
<td>Maximum value for the Hill function for decay rate $\gamma$ of the transient RTF part, which is reached for high doses. For the example plot, the sustained part is set to 0.</td>
<td>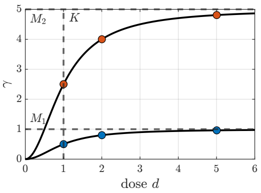<br>$M = 1, 5$<br>$K = 1$<br>$h = 2$</td>
<td>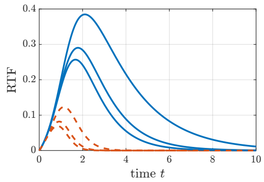<br>$A = 0$<br>$B = 1$<br>$\beta = 1$<br>$\tau = 1$<br>$b = 0$</td>
</tr>
<tr>
<td>$K_\gamma$</td>
<td>The half maximum quantity $K_\gamma$ of the Hill function specifies the dose for which the decay constant $\gamma$ of the transient RTF part is 50% of the maximal rate $M_\gamma$. Roughly speaking, $K_\gamma$ indicates the dose at which the "velocity" is 50% of the maximum velocity of the decay of the transient part.</td>
<td>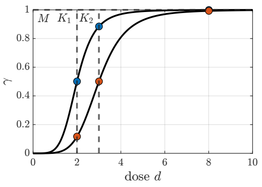<br>$M = 1$<br>$K = 2, 3$<br>$h = 5$</td>
<td>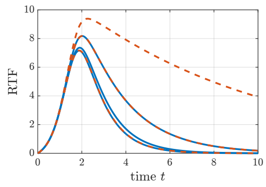<br>$A = 0$<br>$B = 10$<br>$\beta = 10$<br>$\tau = 2$<br>$b = 0$</td>
</tr>
<tr>
<td>$h_\gamma$</td>
<td>The Hill coefficient $h_\gamma$ controls how non-linear the decay constant $\gamma$ depends on the dose. Increasing $h_\gamma$ enhances the sigmoidality of the Hill curve, making it steeper. Large values of $h$ leads to a sudden increase of the "velocity" $\gamma$ for doses close to $K_\gamma$.</td>
<td>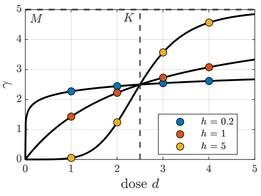<br>$M = 5$<br>$K = 2.5$<br>$h =$<br>$0.2, 1, 5$</td>
<td>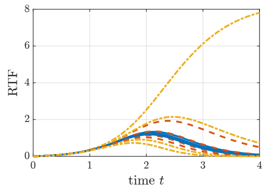<br>$A = 0$<br>$B = 10$<br>$\beta = 1$<br>$\tau = 2$<br>$b = 0$</td>
</tr>
</table>

| | | | | | |
|---|---|---|---|---|---|
| $M_\tau$ | The Hill function of response time $\tau$ is decreasing for increasing doses, with the maximum value $M$ being reached for dose 0. $M_\tau$ corresponds to the time delay that is reached for small doses. For high doses the delay $\tau$ becomes 0 for all $M_\tau$. | 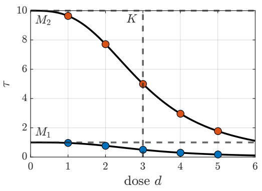  | $M = 1, 10$ <br> $K = 3$ <br> $h = 3$ | 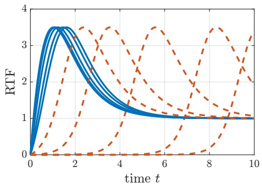  | $A = 1$ <br> $B = 10$ <br> $\alpha = 10$ <br> $\beta = 1$ <br> $\gamma = 1$ <br> $b = 0$ |
| $K_\tau$ | The half maximum quantity $K_\tau$ of the Hill function specifies the dose for which the response time $\tau$ of the transient RTF part is 50% of the maximal response time $M_\tau$. | 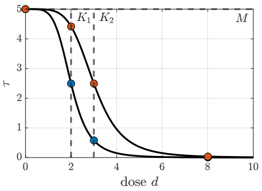  | $M = 5$ <br> $K = 2, 3$ <br> $h = 5$ | 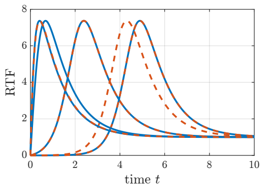  | $A = 1$ <br> $B = 10$ <br> $\alpha = 1$ <br> $\beta = 10$ <br> $\gamma = 1$ <br> $b = 0$ |
| $h_\tau$ | Hill coefficient $h$ for the dose-dependency of the response time $\tau$. Increasing $h_\tau$ enhances the sigmoidality of the Hill curve, making it steeper. Large values of $h$ leads to a sudden decrease of the response time $\tau$ for doses close to $K_\tau$. | 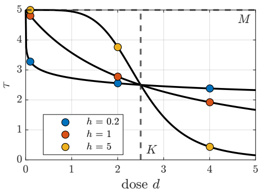  | $M = 5$ <br> $K = 2.5$ <br> $h =$ <br> $0.1, 2, 4$ | 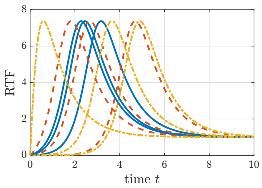  | $A = 1$ <br> $B = 10$ <br> $\alpha = 1$ <br> $\beta = 10$ <br> $\gamma = 1$ <br> $b = 0$ |
| $b$ | Data offset describing a shift along the $y$-axis. For the example plot, $b$ is varied from low ($b = -2$, blue) to high ($b = 3$, red) | The offset $b$ is usually given by a baseline or by a background measurement and is therefore usually not dose-dependent. | | 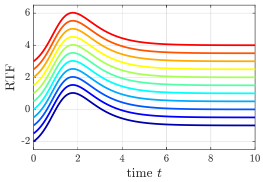  | $A = 1$ <br> $B = 10$ <br> $\alpha = 1$ <br> $\beta = 1$ <br> $\gamma = 1$ <br> $\tau = 1$ |

### S4 Boundaries and Confidence Intervals for all Scenarios

#### S4.1 Modelling doses individually

##### S4.1.1 Dose 1

Table S4: Single dose RTF: Bounds and Confidence intervals for modeled parameters for dose $d = 0\,\mu M$

| Parameter | Lower bound | Estimated value | Upper bound | Confidence interval |
|---|---|---|---|---|
| $A$ | 0.00 | 0.01 | 15.64 | [-Inf, 1.06] |
| $\alpha$ | -0.82 | 0.22 | 0.68 | [-Inf, Inf] |
| $b$ | -5.00 | -5.00 | -1.63 | [-Inf, Inf] |
| $\sigma$ | -0.34 | -0.34 | -0.34 | [NaN, NaN] |
| $\tau$ | -1.92 | 4.79 | 4.79 | [-Inf, Inf] |

##### S4.1.2 Dose 2

Table S5: Single dose RTF: Bounds and Confidence intervals for modeled parameters for dose $d = 1\,\mu M$

| Parameter | Lower bound | Estimated value | Upper bound | Confidence interval |
|---|---|---|---|---|
| $A$ | 0.00 | 3.53 | 15.64 | [2.22, 6.23] |
| $\alpha$ | -0.82 | -0.53 | 0.68 | [-Inf, -0.10] |
| $b$ | -5.00 | -5.00 | -1.63 | [-Inf, Inf] |
| $\sigma$ | -0.34 | -0.34 | -0.34 | [NaN, NaN] |
| $\tau$ | -1.92 | 4.79 | 4.79 | [3.45, Inf] |

##### S4.1.3 Dose 3

Table S6: Single dose RTF: Bounds and Confidence intervals for modeled parameters for dose $d = 2\,\mu M$

| Parameter | Lower bound | Estimated value | Upper bound | Confidence interval |
|---|---|---|---|---|
| $A$ | 0.00 | 4.81 | 15.64 | [4.30, 5.40] |
| $\alpha$ | -0.82 | 0.14 | 0.68 | [-0.22, Inf] |
| $b$ | -5.00 | -5.00 | -1.63 | [-Inf, Inf] |
| $\sigma$ | -0.34 | -0.34 | -0.34 | [NaN, NaN] |
| $\tau$ | -1.92 | 3.71 | 4.79 | [3.04, Inf] |

##### S4.1.4 Dose 4

Table S7: Single dose RTF: Bounds and Confidence intervals for modeled parameters for dose $d = 3\,\mu M$

| Parameter | Lower bound | Estimated value | Upper bound | Confidence interval |
|---|---|---|---|---|
| $A$ | 0.00 | 6.14 | 15.64 | [5.62, 6.72] |
| $\alpha$ | -0.82 | -0.01 | 0.68 | [-0.25, 0.41] |
| $b$ | -5.00 | -5.00 | -1.63 | [-Inf, Inf] |
| $\sigma$ | -0.34 | -0.34 | -0.34 | [NaN, NaN] |
| $\tau$ | -1.92 | 2.97 | 4.79 | [2.50, 3.54] |

#### S4.1.5 Dose 5

Table S8: Single dose RTF: Bounds and Confidence intervals for modeled parameters for dose $d = 4\,\mu M$

| Parameter | Lower bound | Estimated value | Upper bound | Confidence interval |
|---|---|---|---|---|
| $A$ | 0.00 | 5.20 | 15.64 | [4.83, 5.55] |
| $\alpha$ | -0.82 | 0.68 | 0.68 | [-0.01, Inf] |
| $b$ | -5.00 | -5.00 | -1.63 | [-Inf, Inf] |
| $\sigma$ | -0.34 | -0.34 | -0.34 | [NaN, NaN] |
| $\tau$ | -1.92 | 3.40 | 4.79 | [2.49, 3.56] |

#### S4.1.6 Dose 6

Table S9: Single dose RTF: Bounds and Confidence intervals for modeled parameters for dose $d = 5\,\mu M$

| Parameter | Lower bound | Estimated value | Upper bound | Confidence interval |
|---|---|---|---|---|
| $A$ | 0.00 | 6.57 | 15.64 | [6.21, 6.95] |
| $\alpha$ | -0.82 | 0.68 | 0.68 | [0.35, Inf] |
| $b$ | -3.08 | -1.63 | -1.63 | [-Inf, Inf] |
| $\sigma$ | -0.34 | -0.34 | -0.34 | [NaN, NaN] |
| $\tau$ | -1.92 | 3.51 | 4.79 | [3.05, 3.64] |

#### S4.1.7 Dose 7

Table S10: Single dose RTF: Bounds and Confidence intervals for modeled parameters for dose $d = 6\,\mu M$

| Parameter | Lower bound | Estimated value | Upper bound | Confidence interval |
|---|---|---|---|---|
| $A$ | 0.00 | 5.91 | 15.64 | [5.57, 6.27] |
| $\alpha$ | -0.82 | 0.68 | 0.68 | [0.53, Inf] |
| $b$ | -2.50 | -1.63 | -1.63 | [-Inf, Inf] |
| $\sigma$ | -0.34 | -0.34 | -0.34 | [NaN, NaN] |
| $\tau$ | -1.92 | 3.16 | 4.79 | [2.95, 3.29] |

### S4.2 Modelling dose-dependent dynamics

Table S11: Dose Dependencies: Bounds and Confidence intervals for modeled parameters

| Parameter | Lower bound | Estimated value | Upper bound | Confidence interval |
|---|---|---|---|---|
| $K_A$ | -6.00 | -0.07 | 1.78 | [-0.61, 0.20] |
| $K_\alpha$ | -6.00 | 0.38 | 1.78 | [0.09, 0.62] |
| $K_\tau$ | -6.00 | 1.26 | 1.78 | [0.97, Inf] |
| $M_A$ | 0.00 | 7.01 | 15.64 | [5.87, 8.11] |
| $M_\alpha$ | -1.28 | 0.68 | 0.68 | [0.47, Inf] |
| $M_\tau$ | -1.92 | 4.22 | 4.79 | [3.13, Inf] |
| $b$ | -5.00 | -5.00 | 0.89 | [-Inf, Inf] |
| $h_A$ | 0.00 | 0.00 | 1.00 | [-Inf, 0.46] |
| $h_\alpha$ | 0.00 | 0.56 | 1.00 | [0.32, Inf] |
| $h_\tau$ | 0.00 | 0.00 | 1.00 | [-Inf, Inf] |
| $\sigma$ | -3.11 | -0.32 | 0.40 | [NaN, NaN] |

#### S4.3 Modelling condition-dependencies

Table S12: Condition Dependencies: Bounds and Confidence interval for modeled parameters

| Parameter | Lower bound | Estimated value | Upper bound | Confidence interval |
|---|---|---|---|---|
| $K_A$ | -6.00 | -0.37 | 1.78 | [-2.46, 0.04] |
| $K_\alpha$ | -6.00 | 0.56 | 1.78 | [0.44, 0.64] |
| $K_\tau$ | -6.00 | 0.91 | 1.78 | [0.87, Inf] |
| $M_A$ | 0.00 | 6.55 | 15.64 | [5.82, 7.48] |
| $M_\alpha$ | -1.28 | 0.68 | 0.68 | [0.53, Inf] |
| $M_\tau$ | -1.92 | 4.79 | 4.79 | [4.63, Inf] |
| $b$ | -5.00 | -5.00 | 0.89 | [-Inf, -1.15] |
| $\Delta_{K_A}$ | -3.00 | 1.76 | 3.00 | [0.47, Inf] |
| $\Delta_{K_\alpha}$ | -3.00 | -0.24 | 3.00 | [-Inf, Inf] |
| $\Delta_{K_\tau}$ | -3.00 | 0.12 | 3.00 | [-0.98, 0.85] |
| $\Delta_{M_A}$ | -3.00 | 3.00 | 3.00 | [0.82, Inf] |
| $\Delta_{M_\alpha}$ | -3.00 | 1.80 | 3.00 | [0.13, Inf] |
| $\Delta_{M_\tau}$ | -3.00 | 2.00 | 3.00 | [1.35, Inf] |
| $\Delta_{h_A}$ | -3.00 | -0.11 | 3.00 | [-1.17, 0.30] |
| $\Delta_{h_\alpha}$ | -3.00 | 0.59 | 3.00 | [0.14, Inf] |
| $\Delta_{h_\tau}$ | -3.00 | -0.51 | 3.00 | [-Inf, 1.39] |
| $h_A$ | 0.00 | 0.00 | 1.00 | [-Inf, Inf] |
| $h_\alpha$ | 0.00 | 0.40 | 1.00 | [0.28, 0.48] |
| $h_\tau$ | 0.00 | 1.00 | 1.00 | [0.08, Inf] |
| $\sigma$ | -3.11 | -0.36 | 0.40 | [-0.42, -0.30] |
